# Genome-wide identification and characterization of *Solanum tuberosum BiP* genes reveals the role of the promoter architecture in BiP gene diversity

**DOI:** 10.1101/2020.05.16.098244

**Authors:** Venura Herath, Mathieu Gayral, Nirakar Adhikari, Rita Miller, Jeanmarie Verchot

**Affiliations:** Texas A&M Agrilife Center in Dallas, Dallas, TX 77953; Department of Plant Pathology & Microbiology, Institute for Plant Genomics and Biotechnology, Texas A&M University, College Station, TX 77802; Department of Biochemistry and Molecular Biology, Oklahoma State University, Stillwater, OK 77845; Department of Agriculture Biology, Faculty of Agriculture, University of Peradeniya, Sri Lanka

## Abstract

The endoplasmic reticulum (ER) immunoglobulin binding proteins (BiPs) are molecular chaperones involved in normal protein maturation and refolding malformed proteins through the unfolded protein response (UPR). Plants BiPs belong to a multi-gene family contributing to development, immunity, and responses to environmental stresses. This study identified three *BiP* homologues in the *Solanum tuberosum* (potato) genome using phylogenetic, amino acid sequence, 3-D protein modeling and gene structure analysis. These analyses revealed that *StBiP1* and *StBiP2* grouped with *AtBiP2*, whereas *StBiP3* grouped with *AtBiP3*. While the protein sequences and folding structures are highly similar, these *StBiPs* are distinguishable by their expression patterns in different tissues and in response to environmental stressors such as treatment with heat, chemicals, or virus elicitors of UPR. *Ab initio* promoter analysis revealed that potato and Arabidopsis *BiP1* and *BiP2* promoters were highly enriched with cis regulatory elements (CREs) linked to developmental processes, whereas *BiP3* promoters were enriched with stress-related CREs. The frequency and linear distribution of these CREs produced two phylogenetic branches that further resolve the groups identified through gene phylogeny and exon/intron phase analysis. These data reveal that the CRE architecture of *BiP* promoters potentially define their spatio -temporal expression patterns under developmental and stress related cues.

## Introduction

The ER binding immunoglobulin protein (BiP), also known as the glucose receptor protein 78 (GRP78) is conserved across evolutionary kingdoms and is one of the best characterized molecular chaperones in the endoplasmic reticulum (ER). BiP guides the co-translational translocation of nascent proteins into the ER, chaperones protein folding and maturation. BiP plays a key role in protein quality control by identifying and refolding misfolded proteins. BiP directs the post-translational translocation of proteins out of the ER. BiP contains an N-terminal nucleotide-binding domain (NBD) and a C-terminal substrate-binding domain (SBD). The NBD consists of two lobes surrounding the allosteric ATP-binding site, which modulates the substrate-binding activities. SBD consists of two subdomains, SBDβ and SBDα, which enable BiP to bind the hydrophobic surfaces of newly translated proteins to protect them from aggregation in an ATP-dependent manner. The SBDβ is a pocket with two primary loops that surround the nascent polypeptide and, the SBDα lid is covering this pocket^1^. All BiP proteins have C-terminal HDEL or KDEL signaling motif, which ensures their retention and function in the ER and an ATP-binding domain near the N-terminus^2^.

Across eukaryotes, BiPs are among the most abundant chaperones in the ER and are directly engaged in regulating the unfolded protein response (UPR). Under normal conditions in mammalian cells, BiP binds to and inhibits the three ER stress sensors, protein kinase RNA-like ER kinase (PERK), activating transcription factor 6 (ATF6), and inositol-requiring enzyme 1α (IRE1α)^3,4^. BiP releases the sensors to activate ER-to-nucleus signaling cascades. In yeast, Kar2p (BiP orthologue) binds to and inhibits the Ire1p, which is the master regulator of UPR. Dissociation of Kar2p/BiP releases the Ire1p/Ire1α to oligomerize and then splice the mRNA controlling production of the Hac1/XBP1 transcription factor^5^. In Arabidopsis, the ER stress sensors include IRE1a, IRE1b, bZIP28, and bZIP17. The IRE1a and IRE1b splice the mRNA controlling production of the bZIP60 (Hac1p/XBP1 orthologue) transcription factors. The bZIP28 and bZIP17 are transcription factors with transmembrane domains that are cleaved at the ER upon activation. In Arabidopsis, BiP regulates cleavage and removal of the bZIP28 transmembrane domain for nuclear translocation of the transcription factor^6,7^.

In contrast to yeast and mammals, plant *BiP* genes belong to a multi-gene family. *Arabidopsis thaliana* has three well-studied *BiP* proteins. *AtBiP1* and *AtBiP2* are almost identical (99% protein identity) and are expressed ubiquitously^8,9^. *BiP*s are highly expressed in tissues with high levels of cellular secretory activity or rapid cell proliferation. For example, *AtBiP1* and *AtBiP2* expression is well correlated with the synthesis of seed storage proteins and is induced in flower organs and endosperm tissue^9,10^. AtBiPs also mediate fusion of polar nuclei during male and female gametogenesis^11–13^. AtBiP3, which is less conserved (80% protein identity with AtBiP1/2), is shown to be only expressed in response to ER stress^6,8^. The three Arabidopsis *BiP* promoters have UPR-dependent *cis*-regulatory elements (CREs) known as UPRE and ERSE sequences that are transcription targets of bZIP60, bZIP28 and bZIP17^8,14^. The *AtBiP* promoters also have UPR-independent CREs for gene induction by multiple factors having an additive effect on the levels or timing of gene induction^15^.

According to the literature the Arabidopsis, wheat (*Triticum aestivum*), citrus (*Citrus sinensis*), soybean (*Glycine max*), pepper (*Capsicum annuum* L.) and tobacco (*Nicotiana tabacum*) BiPs exhibit protective functions against numerous abiotic stresses including heat, cold, salt, heavy metal and osmotic stresses^16–21^. In soybean and tobacco BY-2 cells, *GmBiPs and AtBiP2* overexpression attenuate ER stress-induced cell death signaling through the N-rich protein (NRP) and vacuolar processing enzyme^1–4^. The Arabidopsis, pepper, rice, soybean, and wheat genomes encode three or four BiPs which have been annotated and, extensive studies have revealed that many of these plant BiP proteins play crucial roles in biotic stress resistance and plant innate immunity^5–9^. Several studies link BiPs to the regulation of ER stress-mediated cell death, hypersensitive cell death, and non-host HR programmed cell death induced by *Xanthomonas oryzae* pv. *Oryzae* in rice, *Pseudomonas syringae* pv. *maculicola* in Arabidopsis, and *Pseudomonas syringae* pv. tomato in soybean and tobacco, *Phytophthora sojae* in soybean^10–14^.

Considering only the *Solanaceae* family of plants, which includes a large number of agronomically important species, the first study that produced partial or complete cDNAs for tobacco BiPs and reported these genes belong to a multigene family was in 1991. This report was before the genome of any *Solanaceae species* was sequenced and annotated. Six genes were identified and classified based on the uniqueness of their 3’ untranslated sequences and were named BiP-like protein (BLP)1, BLP2, BLP3, BLP4, BLP5, and BLP8. Certain BLP proteins were reported to show tissue-specific patterns of expression, stress-related expression, and responsiveness to ER stress induced by tunicamycin treatment ^6,15^. More recent studies have reported the cDNA sequences for three *BiP* homologs in *C. annuum* and *S. lycopersicum* and the annotated loci are publicly available in the genome databases for these plant species. The genome annotation for *N. benthamiana* and *N. tabacum* has been slower to advance. Recent studies have reported the cDNA sequences for four BiP homologs in *N. benthamiana* and *N. tabacum* and named them based on homology with *S. lycopersicum* or *Arabidopsis* BiPs and reported their significant role in biotic defense and plant immune responses ^12,16^. While these transcripts have been identified, many of the loci sequences for these *Nicotiana spp*. are incomplete. Furthermore, the putative transcript or loci IDs provided in databases have significantly changed since the first reported tobacco BiPs making it difficult to cross-reference recently reported sequences with historic information. Although the *Solanaceae* family includes several agriculturally important crops, the *BiP* gene family in *Solanaceae* species remains largely unexplored, poorly annotated, and only partially characterized. This leaves a large gap in our knowledge about one of its major stress response pathways, the UPR.

The potato genome sequence was first published in 2011 and the genome sequence consortium has been actively updating the sequence accessions (solgenomics.net). Potato is the third most important global food crop in terms of human consumption (CIP; www.cipotato.org) and is threatened by many fungal, bacterial, and viral pathogens for which resistance likely requires activities of the BIP gene family. This study identifies and characterizes the *BiP* gene family in potato (*Solanum tuberosum*), that we now know encompasses three BiPs. We analyzed their evolution, structural features, promoter architecture, and their expression under both development and stress-induced conditions. This study provides new insights that suggest their promoter architecture is key to differentiate the various roles of potato *BiP*s in development and during ER stress. Research to discover new genes and to characterize the gene networks associated with the ER stress responses is essential. These findings will be useful to develop strategies to mitigate the detrimental effects of environmental stresses including pathogen attacks which are responsible for significant yield and quality losses in crops worldwide.

## Results

### Identification and phylogenetic analysis of StBiP

To identify BiP homologs in *Solanaceae* (*Capsicum annuum, Nicotiana attenuata, N. benthamiana, N. tabacum, Solanum lycopersicum, S. tuberosum*), we used the Arabidopsis BiP proteins to query the Ensemble Plants and Solanum Genome Network (SGN) using BLASTP ^17,18^. The Arabidopsis BiPs (AtBiP1-3) used for this search were obtained from Ensemble Plants. We also carried out a similar search to retrieve a broader set of putative plant BiP homologs for *Brachypodium distachyon, Glycine max, Oryza sativa, Sorghum bicolor, Triticum aestivum, Zea mays* from Ensemble Plants. We selected proteins that satisfied the e-value of e^-25^ and categorized the retrieved protein sequences as likely functional BiPs based on three predefining criteria: 1) predicted localization to the ER; 2) five internal domains for ATP hydrolysis and substrate binding; 3) ER retention signal (XDEL). We recovered 46 BiPs from both searches and identified three putative *S. tuberosum* BiPs as PGSC0003DMG400012254, PGSC0003DMG400018544, and PGSC0003DMG400024707 (Table 1 and Supplementary Table S1). Here we renamed them *StBiP1, StBiP2*, and *StBiP3* respectively. The amino acid sequence of StBiP1 and StBiP2 are highly similar (92%), whereas StBiP3 is more distantly related to StBiP1 and StBiP2 (75%). The lengths of most BiPs were in the range of 639 to 678 amino acids and their molecular weights were between 71 and 75 kDa. There was one soybean BiP that was 581 amino acids in length and 63.7 kDa and another *N. benthamiana* BiP that was 851 amino acids and 95.8 kDa. The protein PIs were generally between 4.71 and 5.03 (Table 1).

We constructed a phylogenetic tree using PhyML (v. 1.5) with 1000 bootstraps (Fig. 1) in Seaview (http://doua.prabi.fr/software/seaview). This analysis included the 41 plant *BiP*s as well as five vertebrate and *Saccharomyces cerevisiae BiP*s which we retrieved from the corresponding Ensemble databases (Table 1 and Supplementary Table S1)^19^. The phylogenetic tree included three distinct branches representing plants, animals, and fungi, consistent with the evolutionary divergence of BiPs. Among these plant *BiP*s, we observed two distinct groups. Group A consists of ten *BiP*s representing both monocots and dicots and, includes the *AtBiP3* and *StBiP3*. The *AtBiP3* was distinct from the other Group A members. The monocot and *Solanaceae BiP3*-like proteins clustered in separate sub-groups (MIII and SIII). Surprisingly, there were no *B. distachyon, Z. mays, or G. max* genes represented in Group A. These results indicate that there is an *AtBiP3* specific lineage represented in *Solanaceae*.

**Table 1.**
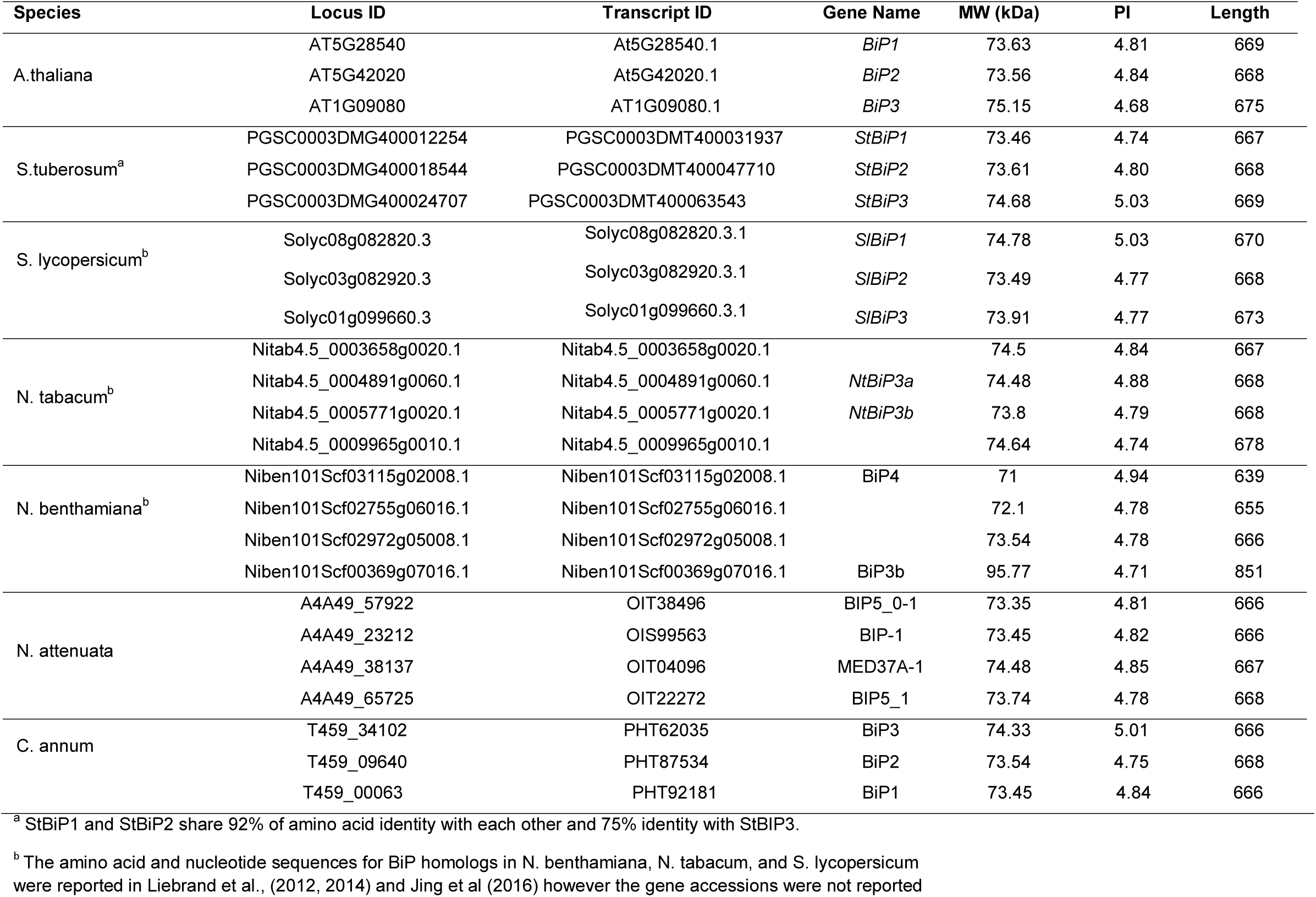
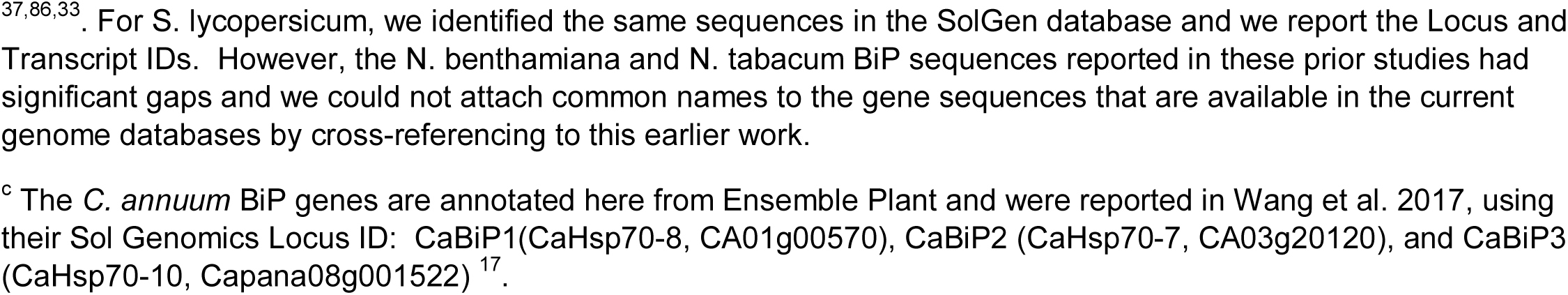
Molecular characterization of *Arabidopsis thaliana* and *Solanaceae* genes.

| Species | Locus ID | Transcript ID | Gene Name | MW (kDa) | PI | Length |
| --- | --- | --- | --- | --- | --- | --- |
| A.thaliana | AT5G28540 | At5G28540.1 | *BiP1* | 73.63 | 4.81 | 669 |
|  | AT5G42020 | At5G42020.1 | *BiP2* | 73.56 | 4.84 | 668 |
|  | AT1G09080 | AT1G09080.1 | *BiP3* | 75.15 | 4.68 | 675 |
| S.tuberosum<sup>a</sup> | PGSC0003DMG400012254 | PGSC0003DMT400031937 | *StBiP1* | 73.46 | 4.74 | 667 |
|  | PGSC0003DMG400018544 | PGSC0003DMT400047710 | *StBiP2* | 73.61 | 4.80 | 668 |
|  | PGSC0003DMG400024707 | PGSC0003DMT400063543 | *StBiP3* | 74.68 | 5.03 | 669 |
| S. lycopersicum<sup>b</sup> | Solyc08g082820.3 | Solyc08g082820.3.1 | *SIBiP1* | 74.78 | 5.03 | 670 |
|  | Solyc03g082920.3 | Solyc03g082920.3.1 | *SIBiP2* | 73.49 | 4.77 | 668 |
|  | Solyc01g099660.3 | Solyc01g099660.3.1 | *SIBiP3* | 73.91 | 4.77 | 673 |
| N. tabacum<sup>b</sup> | Nitab4.5_0003658g0020.1 | Nitab4.5_0003658g0020.1 |  | 74.5 | 4.84 | 667 |
|  | Nitab4.5_0004891g0060.1 | Nitab4.5_0004891g0060.1 | *NtBiP3a* | 74.48 | 4.88 | 668 |
|  | Nitab4.5_0005771g0020.1 | Nitab4.5_0005771g0020.1 | *NtBiP3b* | 73.8 | 4.79 | 668 |
|  | Nitab4.5_0009965g0010.1 | Nitab4.5_0009965g0010.1 |  | 74.64 | 4.74 | 678 |
| N. benthamiana<sup>b</sup> | Niben101Scf03115g02008.1 | Niben101Scf03115g02008.1 | BiP4 | 71 | 4.94 | 639 |
|  | Niben101Scf02755g06016.1 | Niben101Scf02755g06016.1 |  | 72.1 | 4.78 | 655 |
|  | Niben101Scf02972g05008.1 | Niben101Scf02972g05008.1 |  | 73.54 | 4.78 | 666 |
|  | Niben101Scf00369g07016.1 | Niben101Scf00369g07016.1 | BiP3b | 95.77 | 4.71 | 851 |
| N. attenuata | A4A49_57922 | OIT38496 | BIP5_0-1 | 73.35 | 4.81 | 666 |
|  | A4A49_23212 | OIS99563 | BIP-1 | 73.45 | 4.82 | 666 |
|  | A4A49_38137 | OIT04096 | MED37A-1 | 74.48 | 4.85 | 667 |
|  | A4A49_65725 | OIT22272 | BIP5_1 | 73.74 | 4.78 | 668 |
| C. annum | T459_34102 | PHT62035 | BiP3 | 74.33 | 5.01 | 666 |
|  | T459_09640 | PHT87534 | BiP2 | 73.54 | 4.75 | 668 |
|  | T459_00063 | PHT92181 | BiP1 | 73.45 | 4.84 | 666 |

<sup>a</sup> StBiP1 and StBiP2 share 92% of amino acid identity with each other and 75% identity with StBiP3.
<sup>b</sup> The amino acid and nucleotide sequences for BiP homologs in *N. benthamiana*, *N. tabacum*, and *S. lycopersicum* were reported in Liebrand et al., (2012, 2014) and Jing et al (2016) however the gene accessions were not reported
<sup>37,86,33</sup>. For *S. lycopersicum*, we identified the same sequences in the SolGen database and we report the Locus and Transcript IDs. However, the *N. benthamiana* and *N. tabacum* BiP sequences reported in these prior studies had significant gaps and we could not attach common names to the gene sequences that are available in the current genome databases by cross-referencing to this earlier work.
<sup>c</sup> The *C. annuum* BiP genes are annotated here from Ensemble Plant and were reported in Wang et al. 2017, using their Sol Genomics Locus ID: CaBiP1(CaHsp70-8, CA01g00570), CaBiP2 (CaHsp70-7, CA03g20120), and CaBiP3 (CaHsp70-10, Capana08g001522) <sup>17</sup>.

**Figure 1.**
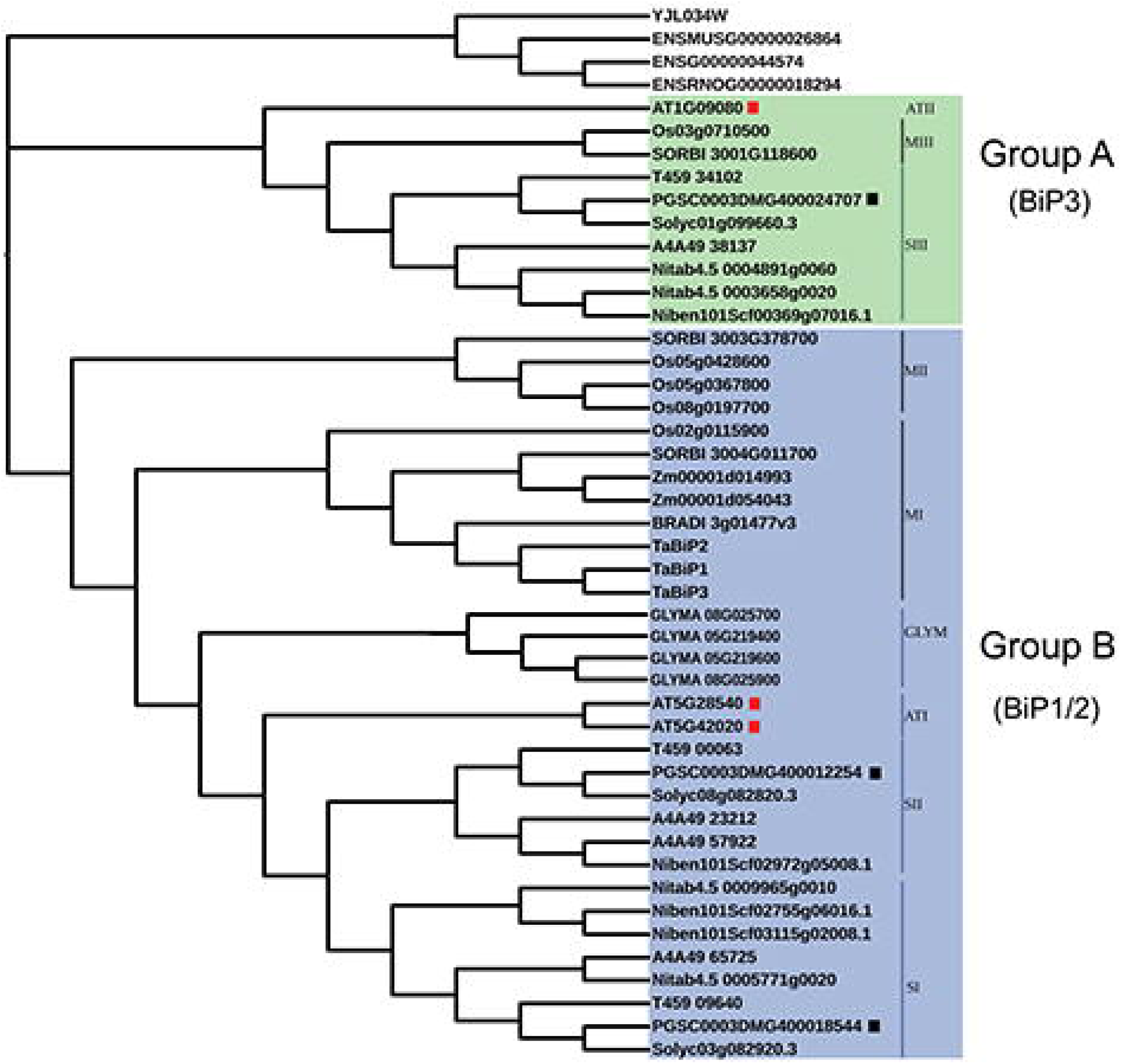
Phylogenetic tree constructed using the PhyML method (v1.5) with Seaview (v4.7) (http://doua.prabi.fr/software/seaview) contains the 46 BiP proteins among fungi, plants, and animals. The output was visualized using iTOL (v4) and the diagrams were compiled, labeled and color coding was added using Adobe Photoshop CC (2017). Group A contains BiPs that cluster with the AtBiP3. The monocot (MIII) and *Solanaceae* (SII) subgroups are identified. Group B is highlighted in blue and represents a larger group that contains AtBiP1 and AtBiP2. The monocots are clustered into two subgroups (MI and MII). Among the dicot BiPs, there are two Solanaceae (SI and SII) subgroups. The soybean and Arabidopsis (GLYM and ATI) BiPs are forming separate branches from the *Solanaceae* BiPs.

Higher diversity was observed among Group B members (Fig. 1). Inside Group B, the monocot and Solanaceae BiPs clustered into two separate subgroups, respectively (MI, MII, SI and SII). In addition, the Arabidopsis and soybean *BiP*s clustered into distinct sub-groups. *StBiP1* and *StBiP2* were present in two *Solanaceae BiP* sub-groups closely related to the sub-group containing the *AtBiP1* and *AtBiP2.* It is worth speculating that Group A has specialized roles in the adaptive evolution of *Solanaceae*.

### Intron/Exon structure of *BiP* genes

The intron positions and the frequency of intron phase combinations in related genes are often interpreted as evidence of a common progenitor. If introns were acquired in a progenitor and stabilized through evolution, then we would expect that the pattern of introns to be non-random. Evidence of random phase distribution of introns suggests that exons were shuffled through evolution which is typically considered as evidence of new functional elements of protein gene products. The position of introns within a codon phase 0,1, or 2, were mapped for each of the 46 *BiP* genes (Fig. 2). The gene sizes of the 46 *BiP*s vary from 2,049 bps to 6,491 bps (Fig. 2). The yeast *BiP* (YJL034W) does not have introns. The representative animal *BiP*s possess seven introns with similar phase patterns suggesting a common origin. Notably, animal *BiPs* have intron phase 2 at the first intron while the plant BIPs used in this study either lack introns or have phase 1 as the first intron. Near the 3’ end is a (1,1) symmetric exon. Among the Group A genes, the *SbBiP3* (SORBI_3001G118600) stands out as unique because it contains one intron and two exons. For all other Group A genes there is central (1,1) symmetric exon surrounded by asymmetric exons in (1,2), (2,1), (1,0) and (0,2) class. The *AtBiP3* (AT1G09080.1) has a (1,2) asymmetric exon and both *AtBiP3* and *OsBiP3* (Os03g0710500) lacked the (0,2) asymmetric exon. The *AtBiP3* had a (1,2) asymmetric exon not found in other Group A genes.

**Figure 2.**
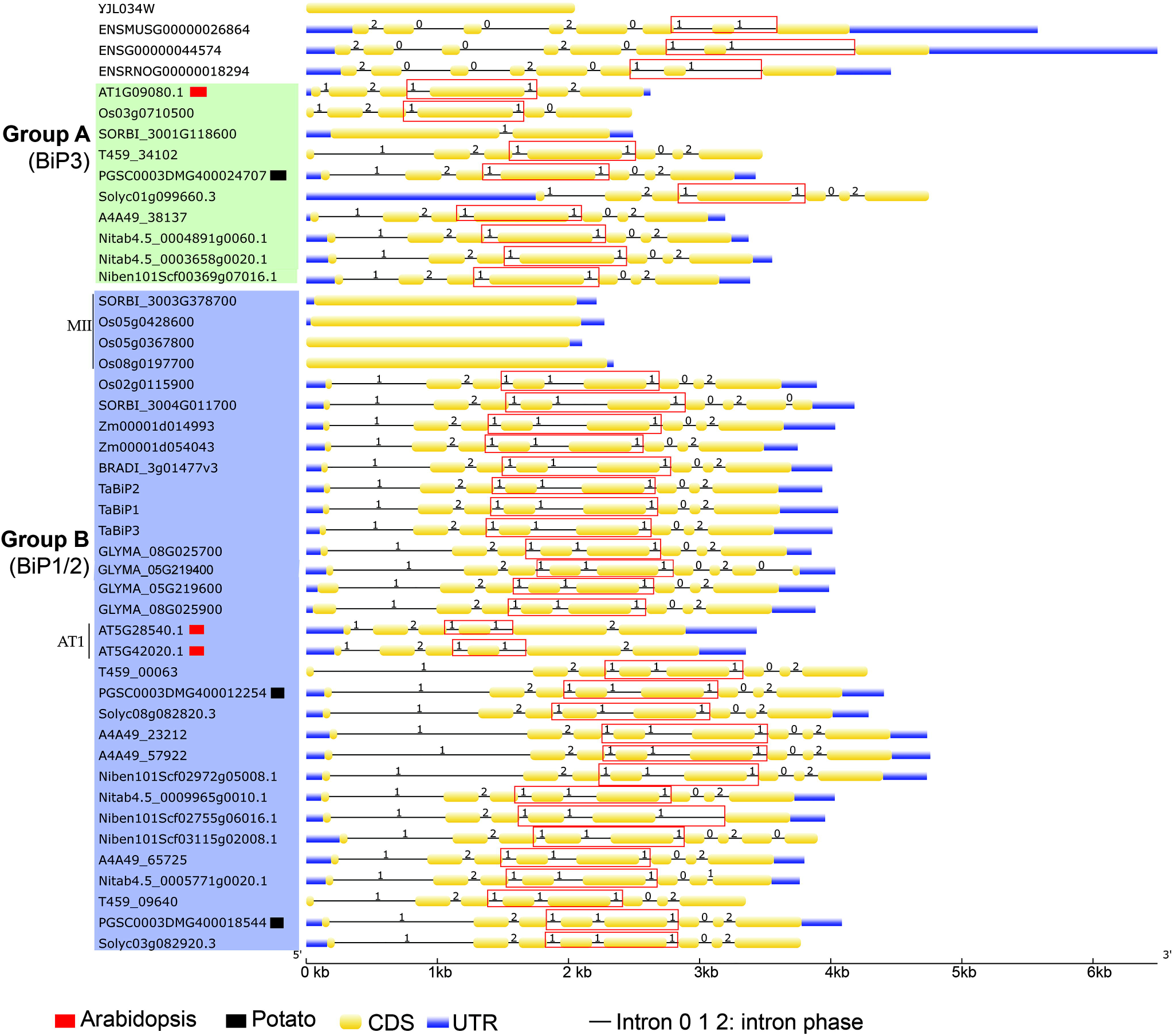
The structures of *BiP* genes, generated using GSDS 2.0 (http://gsds.cbi.pku.edu.cn/), starting from the transcription start site to the end of 3’ untranslated region in fungi, plants, and animals. The gene order is based on Fig. 1 with Group A and Group B BiPs identified in green and blue boxes. The diagram was compiled in Adobe Photoshop (2017) to contain the Group identifiers and colored boxes. The *AtBiP*s and *StBiP*s are marked with red or black squares, respectively, as in Fig 1. The linear representation of each gene includes blue bars indicating 5’ and 3’ untranslated regions, yellow regions representing exons, and lines representing introns. The intron phase 0, 1, 2 are identified above each line. Red boxes surround the (1,1) symmetric exons were added using Adobe Photoshop (2017). Most genes have one or two symmetric exons.

Group B contains a wide assortment of genes including the intron-less rice and sorghum genes (MII subgroup) and all other genes contain between five and eight exons (Fig. 2). The AT1 subgroup contains the *AtBiP1* and *AtBiP2* genes which have the least number of exons (5) and have primarily (1,2), (2,1) (1,2) asymmetric exons and a single (1,1) symmetric exon. The majority of Group B genes have two (1,1,) symmetric exons in the central region, a (1,2) symmetric exon near the 5’ end, and (0,2) or (0.1) asymmetric exon near the 3’ end (Fig. 2). The common intron patterns and phases point towards a common ancestral origin for most of the Group A and Group B BiPs.

### StBiP amino acid sequences and domain structures are highly conserved with AtBiP homologs

It is well known that the N-terminal nucleotide-binding domain (NBD) and C-terminal substrate-binding domain (SBD) of all BiPs are joined by a linker region that is important for allosteric interactions between these two domains ^20,21^. The SBD is further divided into SBDβ and SBDα (Fig. 3a). Because plants have multiple copies of these BiPs, we wanted to know how the structure of these new plant BiPs compared with the canonical structures. To examine this issue, we used the I-TASSER server^22^ to generate three-dimensional structures of AtBiPs and StBiPs using the best available models as threading templates (Fig. 3b). Supplementary Table S2 (see online) provides the confidence score (C-score), the TM-score and the RMSD values to demonstrate the quality of the models for yeast, Arabidopsis and potato BiPs. The topology models were developed using the human BIP and Chinese hamster BiPs as the templates (see Supplementary Table S2 online). TM score and RMSD value correlate with the C-score ^22,23^.

**Figure 3.**
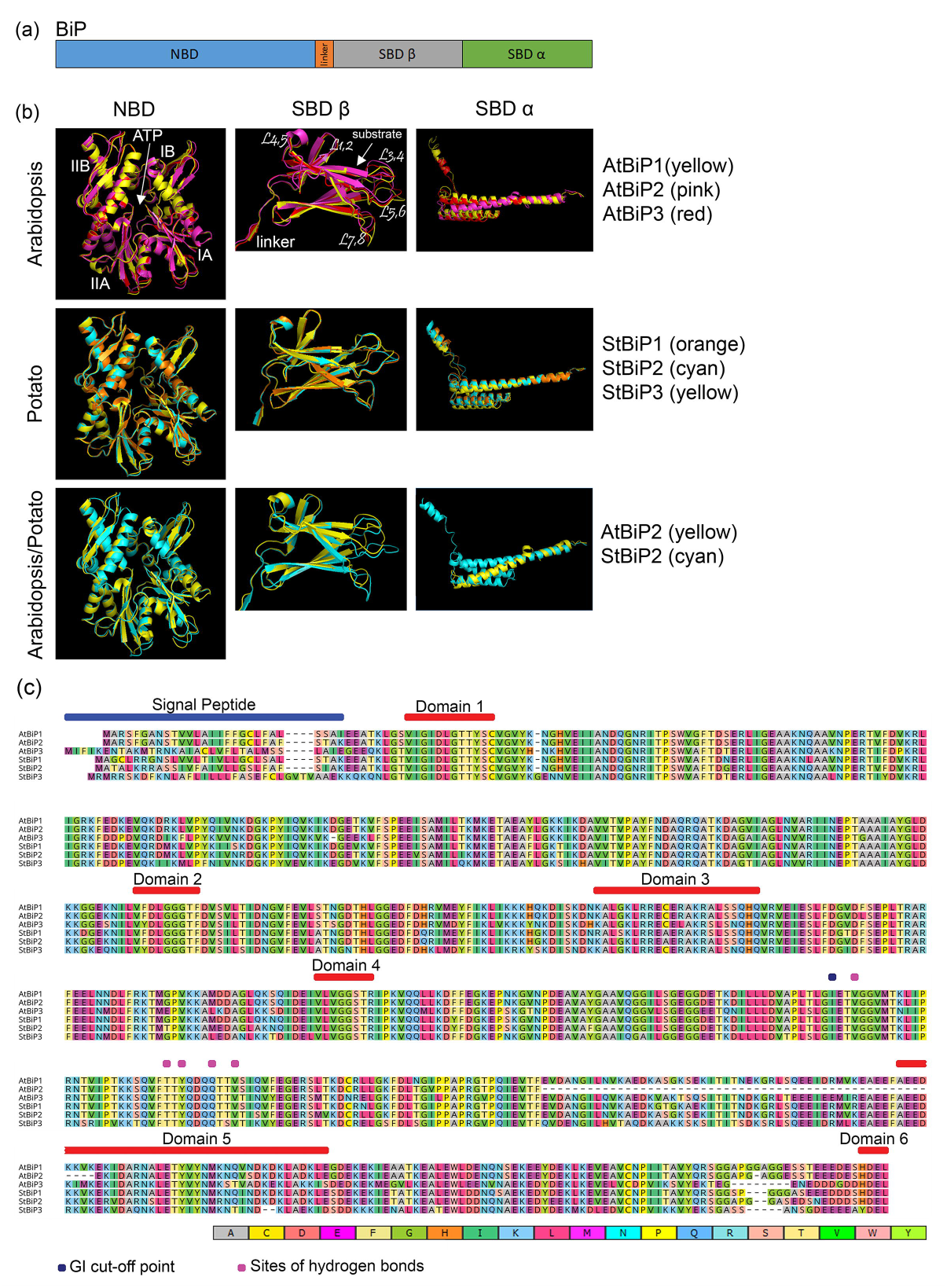
Domain structure of *Arabidopsis* and potato BiPs. a) schematic structures of the BiP domains. b) The 3-D models are superimpositions of NBD, SBDα and SBDβ domains of AtBiPs and StBiPs and were generated using PyMol (v1.7.4) (https://pymol.org/). c) The multiple sequence analysis of the AtBIP and StBiP proteins was generated using MUSCLE built into Seaview (v 4.7) and visualized using Geneious Prime (2019.2.1) The signal peptide, Domain1(β motif), Domain2(γ motif), Domain3(calmodulin-binding site), Domain4 (adenosine-binding motif), Domain5(αβ motif), Domain6(ER retention signal) are identified by bars above the alignment. The identical amino acids are identified by colors pointing to the high conservation among these proteins. The schematic, images and alignments were compiled and labelled using Adobe Photoshop (2017).

Next, we performed 3-D model superimposition, and amino acid sequence alignment of the well-characterized AtBiPs and candidate StBiPs (Fig. 3b, c) by overlaying the NBD, SBDα, and SBDβ of each AtBiP and StBiP to determine if there are observable structural differences between these homologs. There are four structural subdomains oriented around one or more β-sheets (IA, IB, IIA, IIB) that divide the NBD structure into two lobes surrounding a central ATP binding pocket. This general folding structure is highly conserved among the AtBiPs and StBiPs with only minor variation (Fig. 3b). The SBDβ consists of eight β-strands forming a two-layered sandwich with five upper β-sheets and the three lower β-sheets. The structure and amino acid sequences of the SBDβ region of the three AtBiPs and StBiPs are highly conserved. The SBDα consists of four helices. The AtBiP2 has a shorter α-helical domain than AtBiP1 and AtBiP3 that is evident in the 3-D model and corresponds to the gaps in the amino acid alignments that exist upstream of the domain 5 sequence (Fig. 3b, c). The SBDα structure for all AtBiPs and StBiPs, except AtBiP2, are predicted to be highly conserved.

The I-Tasser predicted structure alignments are consistent with the high degree of amino acid conservation seen in the amino acid sequence alignments that were carried out using MUSCLE. The first sequence alignment includes only Arabidopsis and *S. tuberosum* BiPs (Fig. 3c). The second sequence alignment includes all 46 BiPs that were used in the phylogenetic study (see Supplementary Fig. S1 online). Both alignments revealed key conserved amino acid motifs that characterize members of the BiP gene family also occur in the candidate StBiPs (Fig. 3c and Supplementary Fig. S1 online). Within the NBD and SBD are six amino acid sequence motifs that are remarkably well conserved among the 46 BiP sequences. The ATPase binding and hydrolysis functions of BiP require three amino acid domains that are highly conserved, named the β (Domain 1), γ (Domain 2), and adenosine binding (Domain 4). The putative calmodulin-binding motif (Domain 3) located in the ATPase domain is also conserved among the StBiPs. The SBD domain contains the αβ domain (Domain 5), which includes a conserved five-residues substrate-binding core that facilitates hydrogen-bonding with the peptide-substrate backbones, is present in the candidate StBiPs. Finally, an ER retention HDEL signal sequence was present in StBiP1 and StBiP2 while the YDEL signal sequence was present in StBiP3. Thus, these data reveal a high level of conservation among the candidate StBiP1, StBiP2 and StBiP3 proteins and other well-annotated plants BiPs.

### Potato *BiP*s are differentially expressed under developmental and stress conditions

We used the publicly available RNA-seq data that was previously generated from 15 different organs and tissues to analyze the three *StBiP* expression profiles and present the data as heatmaps in Fig. 4a ^24^. Under normal conditions, *StBiP1* and *StBiP2* are constitutively expressed in all tissues and generally show the same pattern of moderate to high expression in stolon, stem, shoot apex, petiole, young and mature tubers. The *StBiP1* is highly expressed in flowers, and the StBiP2 is highly expressed in roots. The *StBiP3* is only expressed in roots under normal growth and development.

**Figure 4.**
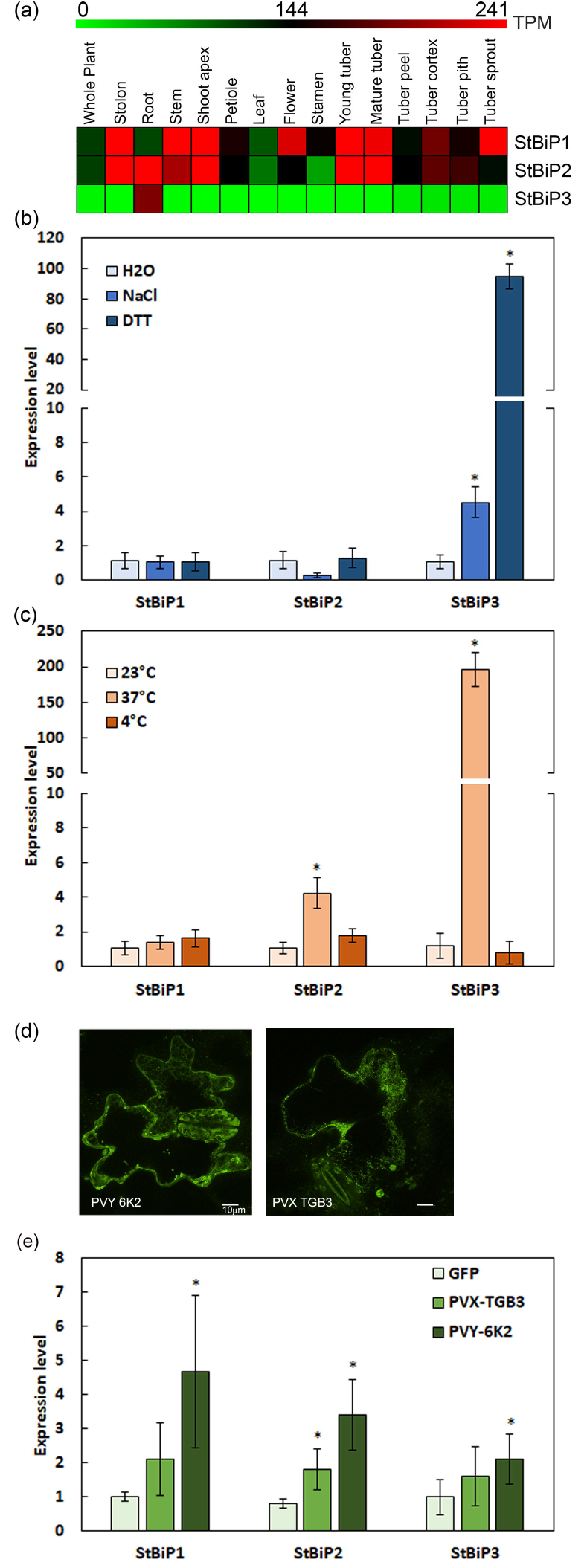
Expression profiles of *StBiP*s under developmental and stress conditions a) a heat map of tissue-specific expression profiles generated from RNA-seq data using MeV (v4.9.0) based on ArrayExpress accession E-MTAB-552^24^ (http://mev.tm4.org/). The transcripts per million (TPM) values of representative transcripts were used to generate the heatmap. The color scale above heatmap shows the expression levels; red indicates high transcript abundance while green indicates low abundance. b-d) The bar graphs were generated using Microsoft Excel (2019) and show the average *StBiP*s transcript levels determined by RT-qPCR following: b) treatment with 2 mM DTT and 150 mM NaCl; (c) heat (37°C) and cold (4°C) treatment; (d) confocal images showing transient expression of GFP fusions with PVY 6K2 and PVX TGB3. (e) transient expression of the potyvirus 6K2 and the potexvirus TGB3 proteins. Abbrev: *significantly different from the control (H_2_O, 23°C and GFP); student t-test; *p*<0.05; n=6)

Since the UPR is an early and rapid response to environmental stress, we investigated whether these genes are engaged to mitigate stress damage caused by abiotic and biotic factors. Extensive prior studies using Arabidopsis, *Nicotiana* species, *Glycine max*, and *Solanum* species treated leaves with DTT or NaCl and carried out time course experiments to demonstrate that the transcriptional response of several UPR related genes including *BiP* can be seen as early as between 1 and 4 h after treatment using RT-qPCR ^25–30^. For simplicity in this study, we infiltrated potato leaves with 2 mM DTT or 150 mM NaCl to induce ER stress and measured BiP transcript levels after 4 h of treatment^30,31^. After four hours, the expression levels of *StBiP1* and *StBiP2* remain unchanged while the *StBiP3* expression in the leaves increased 4.5- and 95-fold, respectively, above the mock control (H_2_O; Fig. 4b).

Prolonged heat and cold treatment are commonly used to measure gene responses to persistent ER stress^28^. Gene expression during persistent ER stress influences plant growth and pro-survival activities. BiP is typically considered to be a core component of the UPR during both early and prolonged responses to ER stress ^32^. Here we exposed potato plants to cold (4 °C), and heat (37 °C) stress for 16 hours^5^. Cold treatment did not lead to changes in expression of any StBiPs (Fig. 4c). In contrast, exposure to heat stress led to a 4-fold increase in *StBiP2* and an approximately 200-fold increase in *StBiP3* in the leaves. Biotic stress was applied by delivering viral proteins that are well known to be specific inducers of the UPR^7,27^. Binary vectors expressing the potato virus X (PVX) TGB3, potato virus Y (PVY) 6K2 proteins fused to GFP, or GFP alone were delivered by agro-infiltration to potato leaves. We confirmed expression of the GFP fusion proteins using confocal microscopy (Fig. 4d). We used RT-qPCR to analyze changes in BiP gene expression after 4 days. As shown in Fig. 4e leaves expressing the PVY-6K2 displayed elevated levels of *StBiP1, StBiP2*, and *StBiP3* transcripts that were between 2.1- and 4.7-fold higher than the GFP controls. Among the leaves expressing the PVX-TGB3, only *StBiP2* was significantly induced. In combination, these data indicate that the expression of *StBiP1* and *StBiP2* are responsive to heat or viral factors. Further, the expression of *StBiP3* is responsive to DTT, NaCl, heat and viral factors that cause ER stress.

### *StBiP* promoters are enriched with developmental and stress-related transcription factor binding sites

To identify the basis of this differential expression, we derived the predicted cis-regulatory elements (CREs) in the three promoters. We selected 1000 bp upstream of the predicted transcription start site for promoter analysis using the New PLACE database (https://www.dna.affrc.go.jp/PLACE/), the TRANSFAC database (release 2019.2 http://genexplain.com/transfac/) and CIS-BP build 2.00 database. We identified CREs belonging to 30 families of transcription factor in the plant *BiP* promoters. All analyzed promoters were enriched with AP2 and bZIP transcription factor binding sites. Then we compared the pattern of CREs in the well characterized *AtBiP*s with *StBiP* promoters in order to further analyze the spatiotemporal regulatory dynamics. The Fig. 5a presents the frequency of the CREs according to the transcription factor families in the *AtBIP* and *StBiP* promoters. Close inspection of these *BIP* promoters revealed the presence of several types of CREs that are well-characterized for their involvement in regulating genes contributing to growth and development, hormone signaling, pathogen defense, and abiotic stress resistance in plants. The basic leucine zipper (bZIP), heat shock transcription factor (HSF), myeloblastosis (MYB) and flowering time regulator SQUAMOSA-PROMOTER BINDING PROTEIN-LIKE (SPL) transcription factor binding sites had the highest representation. Notably, the *StBiP1* lacks an HSF binding site, the *StBiP3* lacks the SPL binding site, which may explain why *StBiP3* is not expressed in vegetative parts of the plant under normal conditions. Cumulatively, there are between 22 and 24 of these sites across all the Arabidopsis and potato *BiP* promoters. These data show that BiPs are regulated by a vast array of transcription factors.

**Figure 5.**
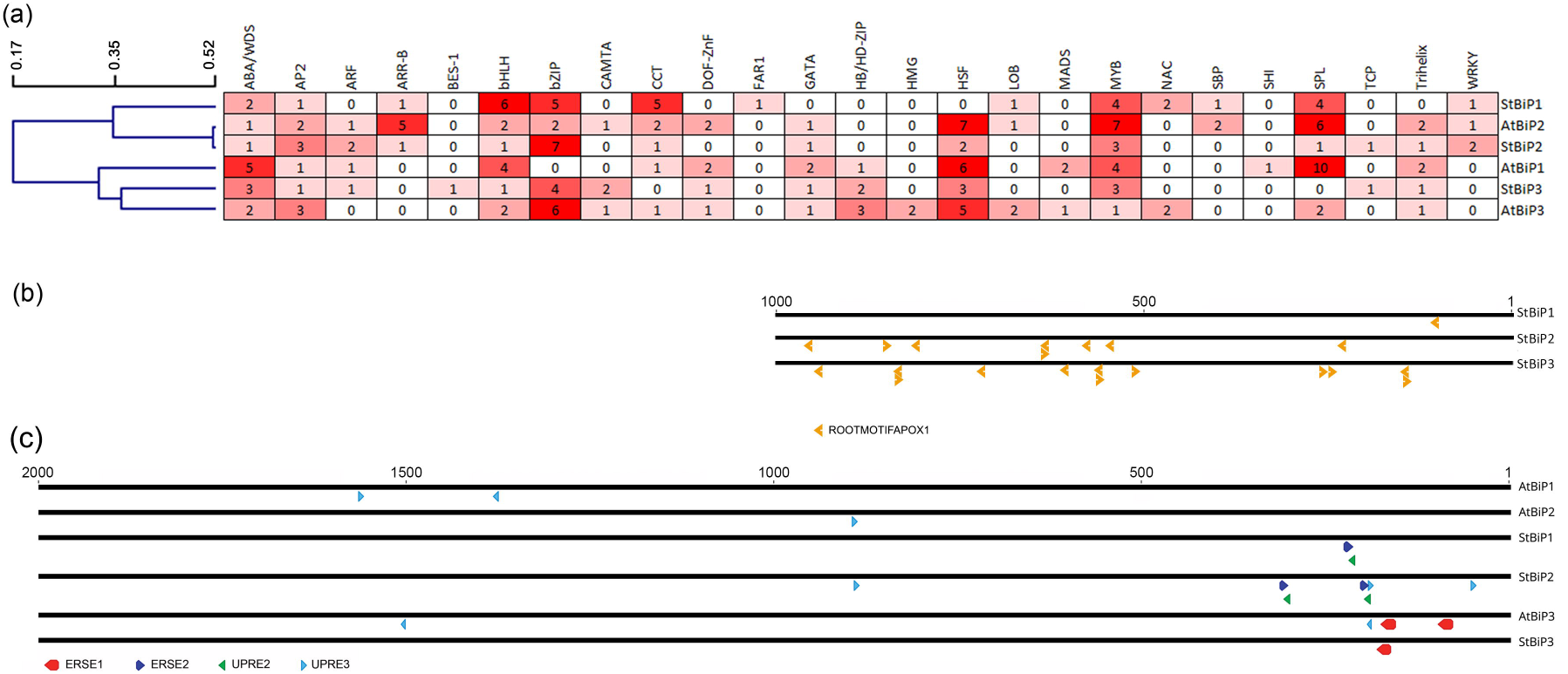
Distribution and frequency of CRE and ER-stress responsive elements on *AtBiP* and *StBiP* promoters. a) The hierarchical clustering of CREs belongs to various transcription factor families identified using the TRANSFAC database release 2019.2 (http://genexplain.com/transfac). The average linkage distance was determined by hierarchical clustering and is presented on the right. The grid provides the number of sites that contain the CREs representing twenty-five transcriptional factor families. Shades of red were used to point to CREs that were highly represented (dark red), moderately represented (medium red), were presented once or twice (pale red), or not represented (white) were identified in *AtBiP* and *StBiP* promoters. The names of the promoters are identified on the right. b) The distribution of ROOTMOTIFTAPOX1 (ATATT) elements (red arrows) in the 1000 bp StBiP promoters are critical for root-preferential expression. These were promoters identified using the New PLACE database (https://www.dna.affrc.go.jp/PLACE). The promoter names are provided on the left. c) The distribution of ER-stress responsive ERSE and UPRE elements in the 2000 bp AtBiP and StBiP promoters. The lines are drawn to scale with the nucleotide position relative to the transcription start site, represented above the lines. The colored arrows represent the positions of the CREs that are engaged by factors in the unfolded protein response. These names of these CRES are identified in the legend. The promoters are named on the right.

The hierarchical clustering of *StBiP*s and *AtBiP*s identified two promoter groups based on the presence and the distribution of CRE on their promoters (Fig. 5a). One cluster consisted of the *StBiP1, StBiP2, and AtBiP2* promoters. The promoter architectures of *AtBiP2* and *StBiP2* were closely related. The second cluster consisted of the *AtBiP1, AtBiP3 and StBiP3* promoters (Fig. 5a). The architecture of the *StBiP3* and *AtBiP3* promoters were also conserved. The promoter architecture of *AtBiP1* and *StBIP1* is distant from the *AtBiP2 and StBiP2*, while the coding region of these two genes are highly conserved.

Given that BiP genes are differentially regulated during growth and developmental processes, Supplementary Table S3 (online) includes specific sets of key transcription factor families and identified CREs that are likely responsible for differential expression of these BiP genes during various biological processes. First, among the highly represented MYB binding sites, the *StBiPs* and *AtBiP1* and *AtBiP2* promoters contain the MYB3R CREs that are components of the DP, Rb-like, E2F, and MuvB (DREAM) complex, pivotal for the onset of cell differentiation processes and organ development ^1,2^. We identified MYB24 CREs in *StBiP1, AtBiP1*, and *AtBiP2* promoters (see Supplementary Table S3 online) which are known to be involved in gibberellic acid and jasmonic acid-mediated stamen development. Interestingly WEREWOLF CREs, which are involved in epidermal cell patterning were present only in the *AtBIP2* promoter. Considering only the potato BiPs, we hypothesize that the presence of such categories of CREs in the *BiP1/2* genes serves to coordinate the elevated levels of *StBiP1* and *StBiP2* as part of the gene network in shoots, stems, stolons, tubers, and stamens^64,65^.

Since these sets of CREs do not explain the differential expression of *StBiPs* in the roots, we further investigated the occurrence of the root-specific ROOTMOTIFTAPOX1 motif (Fig. 5b). This CRE is distributed among all three *StBIPs* with 13 copies in the *StBiP3*, eight copies in the *StBiP2* and only one in the *StBiP1* promoters (Fig 5b). The copy number of root-specific CREs correlates with the respective high, medium, and low levels of *StBiP3, StBiP2* and *StBiP1* in roots seen in Fig. 4.

There are several large families of transcription factors that are well known for their role in protecting key developmental and metabolic processes in response to adverse environments including temperature and water stress. These include the abscisic acid stress and ripening/water deficit stress (ABA/WDS), APETALA2/Ethylene-responsive factor (AP2/ERF), and basic helix-loop-helix (bHLH) transcription factor binding sites. Between 1 and 5 copies of the ABA/WDS and AP2/ERF CREs are present in each AtBiP and StBiP promoters (Fig. 5a). Among the highly represented AP2/ERF binding sites, we noted that the BBM, RAP2.1, B3, and ESR1 are differently represented in each of the *BiP* promoters. For example, the B3 CRE was found only in the *AtBiP1* promoter while ESR1 CRE was only present in the *StBiP1* promoter (Supplementary Table S3). Among the bHLH binding sites, we identified three CREs that are differentially represented. The bHLH28(MYC5) CRE was found on all Arabidopsis and potato promoters except for *StBiP3*. bHLH64 was only present on the *AtBiP3* promoter while BEE2 CREs were found on *StBiP1* and *StBiP3* promoters. These differences suggest fine-tuning of transcriptional activation of the *AtBIP* and *StBiP* genes in response to adverse environmental conditions and/or differential engagement of BiP proteins in cytoprotective processes.

The bZIP transcription factor family in Arabidopsis has 78 members and generally contribute to seed germination, sugar signaling, amino acid metabolism, salt stress, hypo-osmolarity responses, and pathogen defense responses. Regarding sucrose regulation and hormone signaling, the bZIP44/53/GBF6, AS1/OCS/TGA-like, and PEND CREs were prominent in the Arabidopsis and potato *BiP* promoters. Interesting is the occurrence of the bZIP44/bZIP53/GBF6 CREs in StBiP1, StBiP2 and not StBiP3 (see Supplementary Table S3 online). The presence of the as1/OCS/TGA type elements, which are components of regulatory modules associated with hormonal regulation (auxin, salicylic acid, and jasmonic acid) under biotic stresses, wounding, and oxidative stress, might partially explain the activation of all BiP genes by biotic stressors. Only the *AtBiP3* and *StBiP3* promoters had PEND CREs. PEND transcriptional factors are involved in chloroplast DNA replication ^33–35^.

The UPR-related elements were previously identified in the *AtBiP* promoters^36,37^. Here we identify the UPR element II (UPRE-II), UPRE-III, ER stress–response element I (ERSE-I) and ERSE-II in the promoter of *AtBiPs* and *StBiPs* (Fig. 5c, Supplementary Table S3). UPRE-II and ERSE-II elements were overlapping in *StBiP1* and *StBiP2* promoters. The UPRE-III elements were present on *StBiP2* and all *AtBiP* promoters. Only *AtBiP3* and *StBiP3* promoters contain ERSE-I elements that are binding sites for the ER-stress induced AtbZIP28 factor (Fig. 5c)^38,39^. These unique ERSE architectures potentially contribute to the differential expression of BiPs under normal and stressed conditions.

Additional well known CREs with regard to pathogen defense, abiotic stress, and the unfolded protein response are the WRKY and CAMTA binding sites. The WRKY factor binding sites *were present in AtBiP1, StBIP1*, and *StBIP2*, but lacking in the *AtBiP3* and *StBiP3* promoters (Fig. 5A and Supplementary Table S3). WRKY transcription factors modulate plant responses to biotic and abiotic stresses. In particular, WRKY7, −11, and −17 transcription factors are known to control *AtBIP1*/2 expression by suppressing the *AtbZIP28* arm of the UPR during plant immunity ^40^. The *StBIP3* has CAMTA binding sites, which are lacking in the *StBiP1/2* promoters.

## Discussion

The molecular chaperone BiP ensures protein quality control and homeostasis and is an essential regulator of ER stress transducers in development and stress responses. Therefore, the characterization of BiPs is a crucial contribution to the understanding of cellular, tissue-specific and developmentally related responses to ER stress. The *BiP* gene family has been investigated in several plant species, including Arabidopsis, rice, wheat, pepper and citrus^36,37,41–44^. In addition, there are several historic and recent studies using transgenic lines overexpressing various BiPs to characterize their roles in plant tolerance to adverse environmental conditions or demonstrating some evidence for tissue specific regulation^6,15,32,36,43,45^. We have come across many difficulties when we try to retrieve data relevant to the publications especially involving members of the *Solanaceae* family. Some of the difficulties include lack of proper gene nomenclature, annotation, presence of several genome assemblies, and unavailability of relevant nucleotide sequences in databases (Genbank, ENA, etc.)^10,11,15,36^. In order to resolve these issues, we decided to retrieve relevant data directly from curated datasets and updated assemblies while cross comparing with the findings of the above-mentioned studies (Table 1 and Supplementary Table S1). Using the retrieved data, we carried out a comprehensive phylogenetic analysis of both already identified and some novel *BiP*s covering fungi, plant and animal kingdoms in order to identify, characterize BiPs in potato and dissect their evolutionary history.

We have successfully identified three candidate potato *BiP*s (*StBiP1, StBiP2*, and *StBiP3*) in the *S. tuberosum* genome. Further dissection of *StBiP*s using phylogenetic analysis, amino acid sequence analysis, 3-D protein modeling, and gene structure analysis confirmed that these potato gene candidates are true *BiP* genes. The phylogenetic tree divided the *StBiP*s into two subgroups, consistent with previous studies (Fig. 1) ^37,44,46^. Both groups contained monocot and dicot *BiP*s except for Brachypodium. Group A contains *AtBiP*3 and *StBiP*3, while Group B contains the *StBiP*1 and *StBiP*2. Notably, *StBiP*1 and *StBiP*2 seem to have diverged more recently from *AtBiP*1 and *AtBiP*2. Amino acid sequence analysis revealed that *StBiP*1 and *StBiP*2 were highly similar (92%) compared to *StBiP*3 (75%). Similar sequence conservation patterns were observed among the *AtBiP*1, *AtBiP*2 (90%) and *AtBiP*3 (70-76%). Group B *BiP*s show much higher diversification compared to Group A indicating their redundancy and functional diversification. When we examined the six functional domains that define NBDs and SBDs in eukaryotic BiPs, we found that they are highly conserved in the three StBiP proteins. The 3-D model superimposition also revealed a higher degree of structural similarity between the BiP proteins in Arabidopsis and potato. Indicating that their grouping is a result of the variations in non-conserved regions rather than functionally important conserved regions.

Gene structures are reliable indicators of the evolutionary history of gene families^47^. The exon/intron structures and intron numbers vary from zero to eight among the 46 *BiP* genes. Such diversity arises from insertions and deletions during evolution that alters the distance between the start codon and downstream intron peaks. Group A represented by *AtBiP3* is a smaller group whose gene structures and intron phase patterns are conserved except for *AtBiP1*, SORBI_3001G118600, and Os03g0710500. The Group B cluster of *BiP*s has 32 members and excluding the *AtBiP1/2*, SORBI_3004G0117700, Niben101Scf03115g02008.1 and the four *BiPs* that lack introns, all other members have similar intron phase patterns throughout the genes. Although the gene structure and phylogenetic studies differentiated *StBIP1, StBiP2*, and *StBiP3* into different groups, we were unable to assign functional differences for plant development or the management of ER stress based on either of these analyses (gene structure or protein phylogeny) since these differences are not affecting the functional domains of these BiPs. Instead, these data support the functional redundancies often seen in plant experiments where individual *BiP* knockout mutations partially or fully complement other BiP genes in the same plant^27^.

In Arabidopsis, pepper, and rice, the *BiP* genes were primarily distinguishable by their expression profile^42,43,45^. Our results indicate that expression patterns can also distinguish the potato *BiP* genes. Under normal conditions, *StBiP1* and *StBiP2* are expressed in all tissues, whereas *StBiP3* is only expressed in roots. Following the biotic stress induced by the transient expression of the PVX TGB3 and PVY 6K2 proteins, the *StBiP1, StBiP2*, and *StBiP3* showed higher expression in the leaves, which parallels the results reported in Arabidopsis for induction of *AtBiP1, AtBiP2*, and *AtBiP3* ^48^. However, treatments, including high temperature and salt stress, induce only *StBiP3* expression in the leaves. These combined results reveal that *StBiP1* and *StBiP2* involved in specific developmental processes as well as responses to these viral elicitors. The *StBiP3* expresses in the leaves in response to the abiotic stress and the viral elicitors. These functional studies support the phylogenetic protein diversification of StBiP proteins, despite the highly conserved protein structures.

The results presented in this study suggest that the promoter architecture may provide important insights into the various cytoprotective roles of the BiP multi-gene family. This study also demonstrated that a combined analysis, using gene expression profiles and analysis of promoter architecture, is key to differentiating the potato *BiP* homologs. The frequency and the linear distribution of these CREs in the BiP promoter sequences produced two phylogenetic branches that coincide with the separation of Group A (BiP3) and Group B (BiP1/2) in the phylogenetic protein tree. We have identified CRE corresponding to twenty-five transcriptional factor families suggestive of fine regulation of BiPs under various developmental and stress-related stimuli. We propose that the promoter architecture, which controls the gene expression profiles, will be key to investigating the BiPs’ contributions to plant development and adaptation to environmental cues. By understanding the promoter architecture, we can better predict gene functional specificity concerning tissue, timing, and pattern of expression. The distribution and frequency of CREs among *StBiP*s and *AtBiP*s helped us to identify the similarities between the promoter architecture of Group B BiPs. Most interestingly, we were able to differentiate *AtBiP1* and *AtBiP2* based on their promoter variations. They are indistinguishable on gene expression assays due to their high sequence similarity. As a result, identification of their specific functions is a nearly impossible task. However, our results on differential occurrence and distribution of CREs is strongly supporting the hypothesis that they have their own unique functions. Grouping of *StBiP3* and *AtBiP3* strongly supports the conserved regulation dynamics of cellular stress response.

This study successfully identified CREs that are responsible for growth and development, hormonal regulation, and stress response in *StBiP* and *AtBiP* promoters in various frequencies and occurrences. We have identified targets of the DREAM complex that regulate the onset of cell differentiation processes and organ development governed by MYB transcription factors ^49,50^. Also, we identified WEREWOLF CREs (MYB target) that play a major role in epidermal cell patterning ^51–53^). Differential distribution of root-specific ROOTMOTIFTAPOX1 motif was also observed in *StBiP* promoters supporting their differential expression levels in root tissues.

We also recognized CREs representing AP2, bHLH, ABA/EDS that link development, hormonal regulation and stress response in plants. Some of the identified CREs include B3 (mediation of auxin and brassinosteroid dependent development transitions), BBM and RAP2.6 (in cell proliferation, cell morphogenesis, pathogen infection, salt stress, and osmotic stress responses) and RAP2.1 (dehydration response) ^38–40,45,47,48,54^. Most importantly, we have identified various bZIP transcription factor binding CREs important for stress-mediated hormonal signaling in plants. Among the elements, the presence of as1/OCS/TGA (biotics stresses, wounding, and oxidative stress) is validating the role of BiPs in biotic and oxidative stress conditions. Interestingly the distribution dynamics of bZIP binding ER-stress related UPRE and ERSE elements in AtBiP and StBiP promoters have shed light on potential links between ERSE-1 elements and stress-responsive role of AtBiP3 and StBiP3.

We discovered that *StBiP1* and *StBiP2* promoters are highly enriched with CREs related to cell differentiation, organ development, hormone-regulated plant development while the StBiP3 promoter is more enriched with stress-related CREs. However, the presence of pathogen response CRE, AS1/OCS/TGA10 in all three *StBiPs* explain their role in pathogen response. Moreover, AtbZIP17 was already identified in response to virus infection (Gayral et al., 2019; BioRxiv). The StBiP1 and StBiP2 promoters are enriched with UPRE and ERSE elements that can explain their expressions following transient expression of virus proteins. Collectively, the promoter analysis results provide a strong indication of when, where, how and how much each *BiP*s are expressed in potato. Overall, the identification of cis-regulatory architecture will extend our understanding of the complex developmental and stress related *cis*-trans relationships involved in the regulation of *BiP* genes. This provides foundation for dissecting future studies on transcriptional regulation of *BiP* using *in vitro* and *in vivo* strategies including Electrophoretic Mobility Shift Assay (EMSA), Chromatin Immunoprecipitation Sequencing (ChIP-Seq), promoter deletion, and other transgenic approaches. Our findings may ultimately impact not only the functional characterization of *BiP*s but also developing transgene free future-proof crops that can withstand stress conditions using both classical and modern genome editing strategies.

In summary, we identified three BiPs in potato and named them based on their sequence similarity to Arabidopsis BiPs. Their gene structure and domain architectures were highly conserved during the evolution, raising the question of their observed functional diversity under plant growth, development, and stress response. The differences between *BiP* promoter sequences can be used to produce phylogenetic branches that cluster *StBiP2* and *AtBiP2* as well as *StBiP3* and *AtBiP3* into separate subgroups. These promoter differences can also explain the transcriptional differences observed between *BiP1/2* and *BiP3*. Based on our findings, we propose a model-driven by the promoter architecture that initiates spatiotemporal regulation of BiPs under plant growth, development, and stress responses. Accordingly, distribution and frequency of various developmental, stress-responsive and tissue-specific CREs can be considered as the primary determinant of the fine regulation of BiPs both in Arabidopsis and potato.

## Material and Methods

### Genome-wide identification of the *BiP*s in potato and phylogenetic analysis

We used BLASTP and the annotated Arabidopsis *BiP* genes (*AtBiP1*-*3*; AT5G28540.1, AT5G42020.2 and AT1G09080.2) which we retrieved from The Arabidopsis Information Resource (TAIR10) (https://www.arabidopsis.org) to search for BiP homologs in the assemblies of *Brachypodium distachyon* (v3.0), *Capsicum annuum* (ASM51225v2), *Glycine max* (v2.1) *Nicotiana attenuate* (NIATTr2), *Nicotiana benthamiana* (v1.0.1), *Nicotiana tabacum* (v4.5), *Oryza sativa* (IRGSP-1.0), *Sorghum bicolor* (NCBIv3), *Solanum lycopersicum* (SL3.0), *Solanum tuberosum* (in the SolTub_3.0 Genome Assembly), *Triticum aestivum* (IWGSC), *Zea mays* (B73_RefGen_v4), assembly available at the Sol Genomics Network database and the EnsemblPlants database (release 43)^17–19^ within the publicly available EnsemblGenomes project ^55^. An e-value threshold of 10^−25^ was used for the development of the first list of BiPs which included already annotated BiPs as well as candidate sequences of species in *Solanaceae*. Then subcellular localization of the protein sequences of identified potential candidates was assessed using DeepLoc-1.0 ^56^. Only proteins with ER localization signals were identified as putative BiPs. The ENSEMBL Locus IDs were then used to retrieve the gene names and other identifiers from NCBI, UniPro, and TAIR databases (see Supplementary Table S1 online). Multiple sequence analysis was carried out using MUSCLE built into Seaview (v 4.7). The phylogenetic tree was generated using default settings of PhyML (v 1.5) build into Seaview (v 4.7) (http://doua.prabi.fr/software/seaview) with 1000 bootstrap iterations ^57^. The phylogenetic tree was visualized using iTOL (v4) ^58^. Then Adobe Photoshop CC (2017) to compile diagrams into figures and provide labels and add color coding.

### DNA extraction and sequencing

DNA was extracted using rooted cuttings grown in soil inside chambers with a 16 h photoperiod at 22°C for three weeks. Briefly, tissues were harvested in liquid nitrogen and ground using a motor and pestle. DNA extraction and purification were carried out using Mini Genomic DNA Kit (**IBI** Scientific, Peosta, Iowa, USA). DNA quantification was carried out using the Epoch spectrophotometer (Biotek, Winooski, VT) Powerwave XS2 with Gen 5.0 software. PCR amplification of StBiP1, StBiP2, and StBiP3 was carried out using gene-specific primers (Table S4). Platinum™ SuperFi Green DNA Polymerase (Invitrogen) mix was used with PCR amplification conditions as follows; Initial denaturation 98 □C for 2 min, 98 □C 10 sec, 60.9 □C 10 sec, 72 □C for 2 min for 35 cycles, and final extension of 72 □C for 5 mins. Amplified products were visualized using agarose gel (1%) electrophoresis. Sequencing was carried out using ABI 3130 Genetic Analyzer (Applied Biosystems) using sequencing primers (Table S4). The sequencing data of each *StBiP* was analyzed using Geneious Prime® v. 2019.2.1. Gene sequences were deposited at NCBI Genbank under following accession numbers; MN982518 (*StBiP1*), MN982519 (*StBiP2*) and MN982520 (*StBiP3*)

### Gene structures, domain analyses, and protein structure analyses

Intron-exon structures and intron phases of BiPs were visualized using GSDS 2.0 (http://gsds.cbi.pku.edu.cn/)^59^. Conserved domain analysis and visualization were carried out using Geneious Prime (v. 2019.2.1). Images were downloaded from GSDS 2.0. The I-Tasser structure and function prediction tool (Zhang lab.) was used for structural modeling of the protein sequences. The chosen models were analyzed using PyMol (v1.7.4) (https://pymol.org/2/) for observing the protein domains and for the superimposition of the models. All images were downloaded and then constructed into figures using Adobe Photoshop CC (2017).

### Gene expression analysis based on the RNA-seq data

The transcriptomic analysis was carried out using RNA-seq data generated by Potato Genome Sequencing Consortium (PGSC) available at the Expression Atlas Database (https://www.ebi.ac.uk/gxa/home)^24^ generated using RNASeq data coming from ArrayExpress accession number E-MTAB-552. Transcripts per million (TPM) were calculated using raw counts by averaging technical replicates followed by quantile normalization of biological replicates using Limma ^67^. TPMs were used to generate heatmap using MultiExpression Viewer (MeV) version 4.9.0^63^. The diagrams and charts were compiled in Adobe Photoshop CC (2017).

### Plant materials and stress induction

*Solanum tuberosum* cultivar ‘Russet Norkota’ were multiplied by cuttings or *in vitro* on Murashige and Skoog (MS) medium (PhytoTech Labs, Lenexa, KS). Rooted cuttings were grown on soil inside chambers with a 16 h photoperiod at 22°C for three weeks. We subjected vegetative propagated plants in MS media to an overnight 16h of temperature stress by subjecting young sterile plants that are growing in MS media to regular (22°C), high (37°C) or low temperature (4°C). Since the incubators for high and cold temperatures did not have lights, we subjected a control set at regular temperature to an overnight 16 h of dark conditions so plants are exposed to the same light conditions and the only differences between the control and experimental plants are temperature. Leaves were harvested after the 16 h treatment for RNA extraction (below). Experiments requiring infiltrating leaves with solutions of DTT, NaCl or agrobacterium were carried out using rooted plants grown in soil according to Henriquez-Valencia et al. (2015)^30^. For abiotic stress, 2 mM dithiothreitol (DTT), 150 mM sodium chloride (NaCl) or H_2_O as mock treatment were infiltrated into leaves with a 1 mL needle-free syringe. Then leaf samples were harvest at 4 h post infiltration for RNA extraction and RT-qPCR (see below). For viral protein stress, PVX TGB3 and PVY 6K2 sequences were cloned by Gateway Technology (ThermoFisher) in pGWB505 binary vector for produce C-terminal GFP fusions. All plasmids were sequence verified and maintained in *Agrobacterium tumefaciens* sp. strain GV3101. *A. tumefaciens* carrying the GFP expression vector pXF7FNF2.0 was used as a mock control. *A. tumefaciens* harboring TGB3-GFP, 6K2-GFP or GFP were collected and suspended in a solution of 10 mM MES-KOH (pH 5.6), 10 mM MgCl_2_, 200 μM acetosyringone and adjusted to OD_600_=0.7. Then potato leaves (three weeks after rooting) were infiltrated with 1 mL needle-free syringe. These treated leaves were harvest at 4 days post infiltration for RNA extraction (below).

### RNA extraction and RT-qPCR

Leaves were harvested, ground in liquid nitrogen, and RNA was extracted using the RNeasy Plant Mini Kit (Qiagen, Germantown, MD). Total RNA (2µg) was reverse-transcribed using a High-capacity cDNA Reverse Transcription Kit (Applied Biosystems, Foster City, CA) and random primers were employed for cDNA synthesis. After 500-time dilution, qPCRs were performed using the SYBR Green Master Mix (Applied Biosystems), QuantStudio 6 (Thermo Scientific) and specific primers. For detecting *StBiP1* we used the forward primer (GAGGGTGGAGACGAAACTAAAG) and reverse primer (GGGACTTCTTGGTTGGGATAA). For detecting *StBiP2* we used the forward primer (GAGGGTGGAGACGAAACTAAAG) and reverse primer (GCCTTCTTGACAGGAGTCATT). For detecting *StBiP3* we used the forward primer (CTAGTGACAAGTCGCGTCTAA) and reverse primer (GAGCATCCACCTTCTCCTTTAC). For internal control, we used primers that detect *StACTIN58* (PGSC0003DMG400023429). The *StActin58* forward primer (ACCATCGGTGCTGAACGATT) and the reverse primer (TCTATCGGCAATACCGGGGA) were used. The ΔΔCt method was used for calculating relative gene expression. The data represent the mean of two technical replicates for three biological repetitions. Data was analyzed and charted using Microsoft Excel (2019). Charts were compiled using Adobe Photoshop CC (2017).

### Confocal Microscopy to confirm expression of viral proteins fused to GFP

Leaf segments were cut with a scalpel and places on a microscope slide and cover slip and then examined with an Olympus Fluoview FV1000 confocal laser scanning microscope and a 60 X objective (Olympus America Inc. Center Valley, PA). Laser excitation wavelength of 515 nm and Z-dimensions of 5 µm with 0.5 µm/slice. Image J 1.52p software^68^ was used for stacking Z-series images.

### *Ab initio* promoter analysis

Promoter sequences representing 2000 bp from the transcription start site were retrieved from the Ensemble Plants database (http://plants.ensembl.org/; Ensembl Plants release 43, April 2019). CREs were identified using the New PLACE database (https://www.dna.affrc.go.jp/PLACE/), the TRANSFAC database (release 2019.2 http://genexplain.com/transfac/) and the Catalog of Inferred Sequence Binding Proteins (CIS-BP) Database (v 2.00) (http://cisbp.ccbr.utoronto.ca/) on 1000 bp from the transcription start site ^60–62^. Hierarchical clustering was carried out using the Manhattan correlation with the average linkage method in MultiExpression Viewer (MeV)(v4.9.0) http://mev.tm4.org/) ^63^. The output of the hierarchical clustering was retrieved and the heatmap was recreated using Microsoft Excel (2019). UPRE and ERSE elements were identified in the 2000 bp promoter sequence (see Supplementary Table S3 online). Development and tissue-specific expression profiles were obtained from the Expression Atlas Database (https://www.ebi.ac.uk/gxa/home)^24^. Geneious Prime (v. 2019.2.1) was used to annotate and visualize the CREs and the diagrams were compiled using Adobe Photoshop CC (2017).

## Acknowledgments

This work was supported by a grant from NSF (IOS #1759034). Additional support came from the Oklahoma Agriculture Experiment Station (OKL02961) to Rita K Miller and Texas A&M Agrilife to Jeanmarie Verchot

