## Supplementary figures and images for "Genome-wide identification and characterization of *Solanum tuberosum BiP* genes reveals the role of the promoter architecture in BiP gene diversity"

### Supplemental Figure

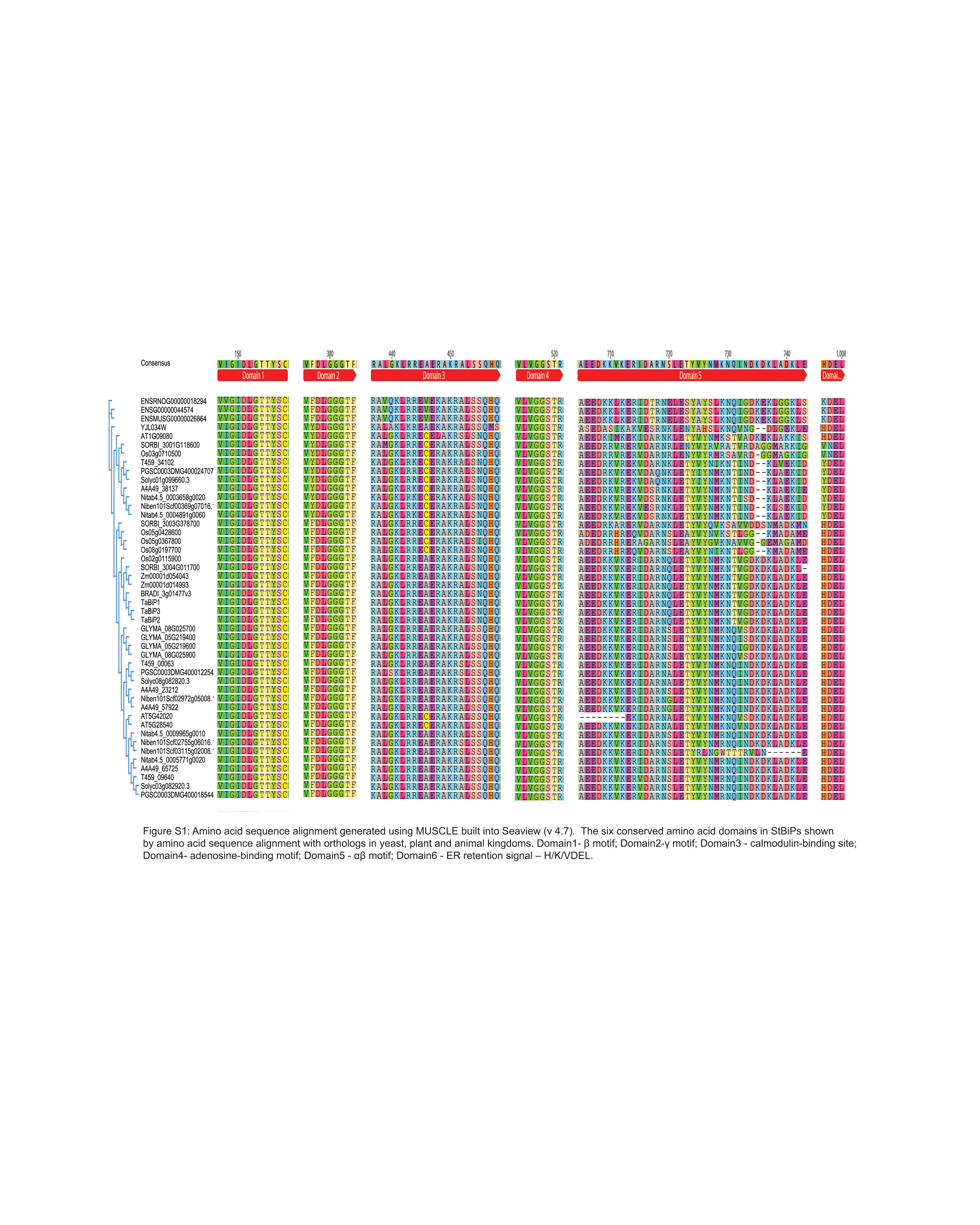
